# In vivo imaging of ferroptosis through nanodynamic changes in lipid membranes

**DOI:** 10.1101/2024.10.25.620222

**Authors:** Ali Yasin Sonay, Benedict Edward Mc Larney, Elana Apfelbaum, Jan Grimm

## Abstract

Ferroptosis emerged as a cell death modality for drug resistant cancer cells, but there are currently no available biomarkers for imaging ferroptosis based therapies. To address this gab, we evaluated the nanodynamic changes in lipid membranes occurring during cell death to explore potential targeting opportunities to image cell death. We nano-sized gaps at late stages of ferroptosis can serve as entry points for dyes that can bind to cellular structures. These changes were accompanied with cellular signaling components similar to platelet activation, with phosphatidyl serine emerging on the surface of the cells and therefore as a potential target for imaging of programed cell death, including ferroptosis. Taking advantage of these changes in cell membrane dynamics, we employed a novel tumor-seeking dye CJ215 that can label apoptotic cells as recently described by us. We show that CJ215 accumulates in ferroptotic cells both in vitro and in vivo by binding to phosphatidyl serine, a process that is prevented by inhibition of ferroptosis. Since phosphatidyl serine exposure also occurs during apoptosis, CJ215 can serve to image both apoptosis and ferroptosis based therapy.

## Main

Ferroptosis is a more recently discovered cell death mechanism that is caused by disruption in cellular redox regulations that ultimately results in excess lipid peroxide accumulation in cellular membranes with subsequent membrane instability and disruptions ^1^. Lipid peroxidation is thought to occur mostly through the Fenton reaction between hydrogen peroxide and ferrous iron, generating hydroxyl radicals that oxidize the polyunsaturated fatty acids in the membrane ^2^. GPX4 is a crucial enzyme that can reduce these lipid peroxides, so either direct inhibition of GPX4^3^ or depletion of its substrate Glutathione (GSH)^4–6^ can lead to ferroptosis. Over the years, several ferroptosis inducers have been discovered such as cystine uptake inhibitor erastin^7^, GPX4 inhibitor RSL3^8^, GSH synthesis inhibitor BSO^9^ or clinically approved iron oxide nanoparticles to name a few^10^. It is widely accepted that significant changes in oxidized lipid species occur during (and are a hallmark of) ferroptosis, which are typically characterized by lipidomic approaches such as measuring lipid peroxidation^11^. Beyond the changes in lipid content, lipids’ localization and membrane integrity are essential factors that maintain membrane homeostasis^12^. This homeostasis emphasizes the boundaries between inside and outside of a cell that in turn are responsible for compartmentalization of cellular processes to control pH, ion levels, membrane potential, metabolites, proteins and enzymatic processes^13^. Dysfunctional antioxidant pathways and lipid peroxidation in the membrane initiates signaling cascades that contribute to ferroptosis, but while intense focus has been given to the lipid species that define ferroptotic cell states^14^, little attention is given to the nature of the membrane damage and lipid exposure during the course of late ferroptosis. In this study, we have investigated the causes and characteristics of nano-sized gaps on the membrane as the cells undergo ferroptosis. Furthermore, we took advantage of these nanodynamic changes to employ a labeling strategy for in vivo monitoring of ferroptosis-based therapy.

### Shared components between platelet activation and ferroptosis

Ferroptosis leads to characteristic membrane swelling and eventual rupture^15^ but the timescales for these events are dependent on cell type, as well as the chosen ferroptosis inducer and therefore vary widely. To derive a more universal ferroptosis framework, we looked at early stage (within 6 hours of erastin treatment) and a late stage (between 12-18 hours post erastin addition) induced ferroptosis. To date, membrane integrity during ferroptosis is analyzed via cell-impermeable dyes that were typically used for apoptosis assays^16^. While this approach reveals membrane damage with high sensitivity, it does not report the size of any generated membrane disruptions during cell death. To overcome this, we employed a 1 nm sized organic dye (DAPI) as well as water soluble quantum dots with size distributions around 5, 10 and 15 nm, respectively **(Supp Fig. 1a)**. Since increasing quantum dot size leads to their red shifted emission spectra, we were able to distinguish passive cellular uptake of different sizes^17^ using flow cytometry during early and late ferroptosis stages (see method section for more details) **(Fig. 1a-b)**. Our results revealed that during early ferroptosis, membrane damage only permitted the uptake of the 1 nm sized organic dye **(Figure 1c)**, whereas during late stage ferroptosis, 1 nm dyes have more pronounced uptake, 5 nm sized QD450 penetrate the damaged membrane but larger quantum dots do not **(Figure 1d)**. Previous studies discussed the signaling aspect of lipid peroxide species and their effect on cell rupture and immune cell activation^18^. We investigated whether membrane swelling and integrity disruption in ferroptosis leads to release of intracellular components to the surrounding cells similar to immunogenic cell deaths such as pyroptosis^19^. We tested different adrenergic, prostaglandin and thromboxane receptor modulators^20^ which are involved in immunogenic responses **(Supp Fig. 2a)** to see their effect on and possible role in cells undergoing ferroptosis. We showed that the thromboxane A2 (TXA2) receptor inhibitor seratrodast rescues the cells from erastin and RSL3 induced ferroptosis whereas other modulators do not greatly influence ferroptosis **(Fig 1e, Supp Fig. 2b)**, Previous studies also showed that inhibition of thromboxane A2 receptor, a G-protein coupled receptor involved in platelet activation, inhibits neuronal ferroptosis by decreasing lipid peroxidation and inhibiting JNK phosphorylation^21^. We reasoned that there are shared components of the ferroptosis and platelet activation pathways such as PIP2-PLC-IP3 receptor-PKC axis^22^ **(Figure 1f)** since seratrodast is a platelet activation inhibitor and platelet activation is also accompanied by similar membrane swelling.

**Figure 1.**
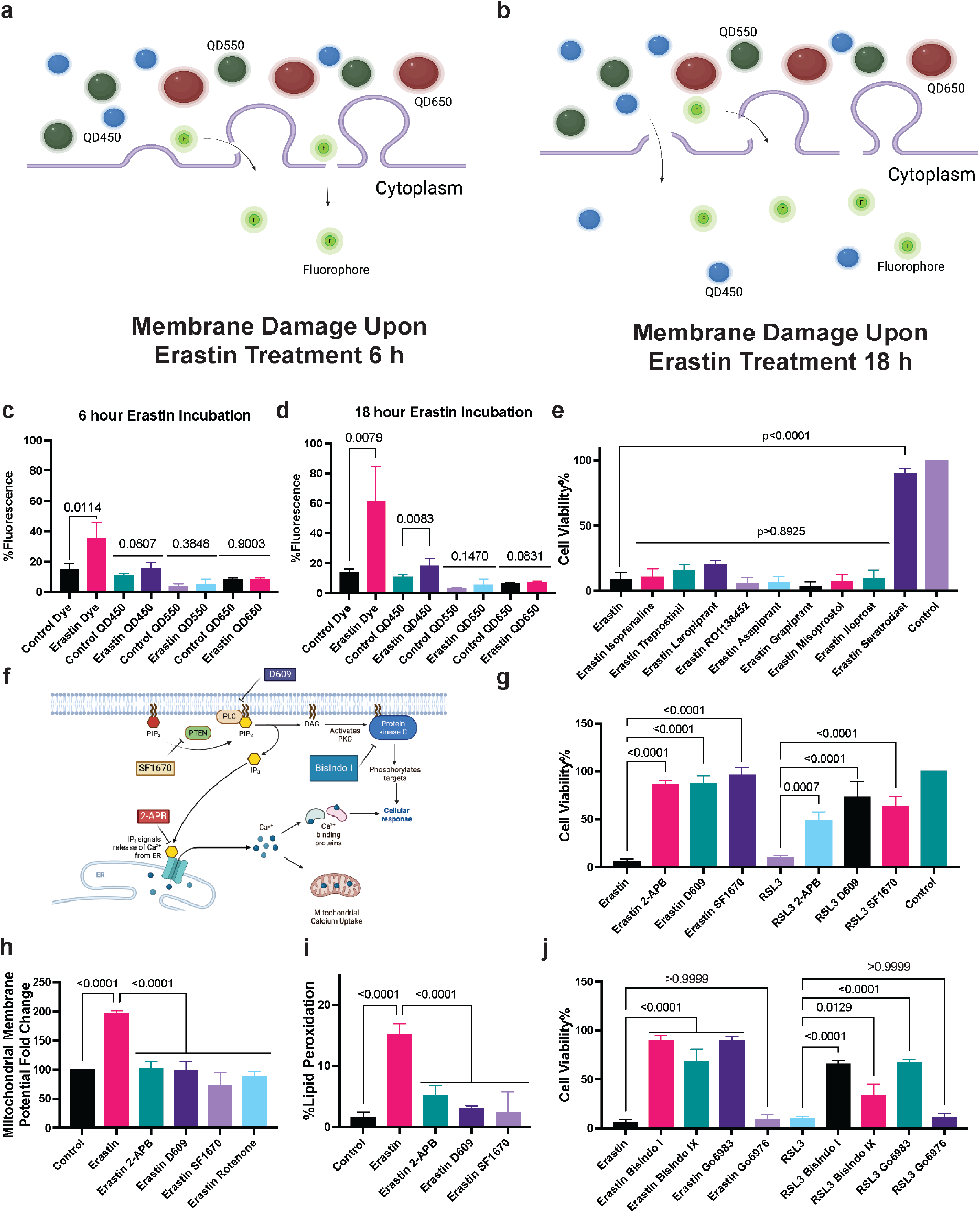
Ferroptosis leads to loss of membrane integrity and shares similar components with platelet activation. **a)** Figure showing loss of cellular membrane integrity showing nano-sized gaps less than 5 nm only allowing dye uptake **b)** Figure showing loss of cellular membrane integrity showing nano-sized gaps less than 10 nm only allowing dye and QD450 uptake, **c)** Cell membrane penetration of different sizes of quantum dots and DAPI upon Erastin treatment for 6 hours showing dye uptake but not quantum dot uptake, **d)** Cell membrane penetration of different sizes of quantum dots and DAPI upon Erastin treatment for 18 hours showing increased uptake of DAPI and 5-8 nm quantum dot with 450 nm excitation **e)** Effect of prostaglandin and thromboxane modulators on erastin induced cytotoxicity showing platelet activation inhibitor seratrodast rescues the cells from ferroptosis. **f)** Figure showing key components of platelet activation where inhibitors of each component are shown in rectangular boxes **g)** IP3R, PLC and PTEN inhibitors 2-APB (2 μM), D609 (5 μM), SF1670 (0.2 μM) respectively rescue the cells from both erastin and RSL3 induced ferroptosis **h)** Erastin induced increases mitochondrial membrane potential measured with JC10 dye which can be inhibited with IP3R, PLC, PTEN inhibitors as well as Electron Transport Chain inhibitor Rotenone (1 μM) **i)** IP3R, PLC, PTEN inhibitors rescues the cells from erastin induced lipid peroxidation, **j)** Role of different PKC inhibitors and their effect on erastin and RSL3 induced ferroptosis. For cytotoxicity assays n=3 biological replicates, for the lipid peroxidation and mitochondrial membrane potential assays n=6 pooled from n=3 independent experiments. Mean ± s.d. p-values are reported above the lines (Ordinary one-way ANOVA with Tukey’s multiple comparisons).

One of the key events during platelet activation is release of calcium from endoplasmic reticulum using IP3 receptors as calcium channels. Our results revealed IP3 receptor inhibitor 2-APB led to the inhibition of both erastin and RSL3 induced ferroptosis. Therefore, we tested next whether IP3 signaling molecule synthesis also influences ferroptosis. Given IP3 is produced from PIP2 signaling molecules via Phospholipase C (PLC), we tested whether inhibition of PLC activity would rescue the cells from ferroptosis. Furthermore, PIP2 levels are tightly regulated by the balance of IP3 kinase (which converts PIP2 to PIP3) and PTEN which converts PIP3 to PIP2. We reasoned inhibition of PTEN would lead to a depletion of PIP2 pool and would prevent cells to undergo ferroptosis. Our results showed both PLC inhibitor D609, PTEN inhibitor SF-1670 rescue the cells from both erastin and RSL3 induced ferroptosis **(Figure 1g)**. We also tested the role of these inhibitors in HT1080 cells and showed that PLC and PTEN inhibitors rescue the cells from erastin and RSL3, while the IP3 receptor inhibition only prevents erastin induced ferroptosis **(Supp Fig. 3a)**. Previous studies also showed erastin treatment led to an increase in mitochondrial membrane potential which is inhibited by electron transport chain inhibitor rotenone^23^, and we demonstrated IP3 receptor, PLC and PTEN inhibitors also decrease the mitochondrial membrane potential similar to rotenone **(Figure 1h)**. Erastin-induced lipid peroxidation can also be prevented using IP3 receptor, PLC and PTEN inhibitors **(Figure 1i)**. Upon PLC activity, PIP2 molecules are hydrolyzed to IP3 and DAG. As a result, both and IP3 receptor induced Calcium release and DAG activity lead to Protein Kinase C (PKC) activation. Yet, there are different classes of PKCs and each have diverse functions depending on the cell type and environmental conditions. Given the redundancy of PKC activation, it is hard to distinguish which PKC isoform is responsible for ferroptosis, therefore we created a small PKC inhibitor library to test multiple PKC inhibitors (see **Supp Fig. 3b for a list of different PKC inhibitors and their targets)** to distinguish whether we can identify the responsible PKC types. We observed that erastin and RSL3 induced ferroptosis could be rescued by the PKC inhibitors BisIndolmaleimide I, BisIndolmaleimide IX, Go6983, but not by Go6976 in MDA MB 435 cells **(Figure 1j)**. In contrast, HT1080 cells treated with erastin can also be rescued by BisIndolmaleimide I, BisIndolmaleimide IX, Go6983, yet RSL3 induced can only be rescued with BisIndolmaleimide I and Go6983 **(Supp Fig. 3c)**. As we systematically assessed the specificity of each inhibitor on PKC isoforms, our results indicated both PKCβII and PKC-γ are involved with ferroptosis depending on different cell lines and ferroptosis inducers which is in accordance with previous studies^24^

### Membrane nanodynamics as potential biomarkers for ferroptosis

As we found shared pathways between platelet activation and ferroptosis such as TXA2 receptor activation and PIP2-PLC-IP3 receptor and PKC pathway, we asked whether phospatidyl serine exposure as a hallmark of platelet activation^25^ also occurs during ferroptotic cell death. This change would not be detected with existing lipidomics approaches as they can only detect the changes in lipid content, not the localization or accessibility of lipids in the membrane. Phosphatidyl serine exposure is typically monitored using Annexin V protein conjugated to an organic dye, which would bind to the outer membrane in apoptotic cells. The Annexin V protein conjugated with a dye is used in conjunction with a membrane impermeable dye that tests membrane integrity in cells undergoing cell death. In the case of apoptosis, phosphatidyl serine exposure occurs when the membrane is intact, so apoptotic cells only show an increase in Annexin V staining without an increase in the membrane impermeable dye. In contrast, as we have shown, damaged membranes with nanosized gaps lead to an increased uptake of membrane impermeable dyes within the early stages of ferroptosis and continue to increase towards the late stages. When we stained early and late stage ferroptotic cells with Annexin V stain, we observed an increase in signal during the late stage erastin-induced ferroptosis which can be inhibited with ferroptosis inhibitor liproxstatin. Similarly, post 2-hours of RSL3 treatment resulted in an increase in Annexin V staining that is rescued by liproxstatin **(Figure 2a)**. Since this increase in Annexin V staining occurred in the late stages of ferroptosis, this change can result from i) exposure of phosphatidyl serine to the outer leaflets and ii) Annevin V-dye conjugate passing through the 5 nm sized gaps and binding to phosphatidyl serine molecules on the inner leaflet. As these mechanisms are not mutually exclusive, both can play a role to label phosphatidyl serine during the late stages of ferroptosis.

**Figure 2.**
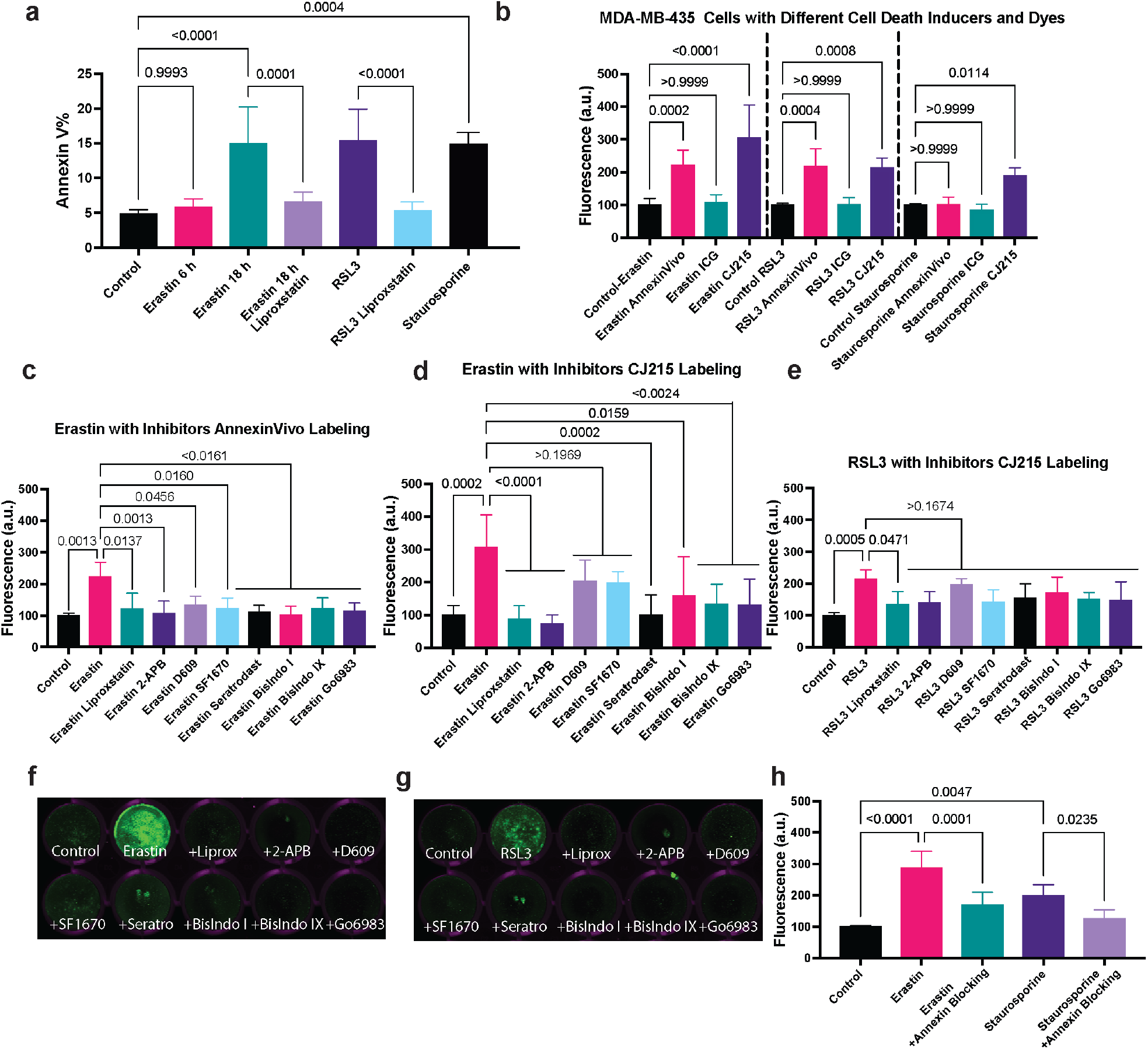
Ferroptosis induction leads to Phosphatidyl serine exposure and can be detected with commercially available dyes Annexin Vivo and CJ215. **a)** Annexin V staining of cells treated with Erastin (15 μM) for 6 and 18 hours and RSL3 (1 μM) treated for 2 hours either with and without Liproxstatin (2 μM) shows an increased fluorescence upon ferroptosis induction which can be decreased with liproxstatin. 18 hour Staurosporine treatment is used as a positive control, **b)** Measuring the fluorescence levels in MDA MB 435 cells treated with erastin, RSL3 and staurosporine using three different dyes AnnexinVivo, ICG and CJ215 showing the effectiveness of CJ215 over the other contrast agents, **c)** Measurement of fluorescence upon 24 hours of erastin treatment (10 μM) along with ferroptosis inhibitors using AnnexinVivo staining **d)** Measurement of fluorescence upon 24 hours of erastin treatment (10 μM) along with ferroptosis inhibitors using CJ215 staining **e)** Measurement of fluorescence upon 6 hours of RSL3 treatment (1 μM) along with ferroptosis inhibitors using CJ215 staining **f)** Representative image of erastin treated cells with and without ferroptosis inhibitors using CJ215 staining **g)** Representative image of RSL3 treated cells with and without ferroptosis inhibitors using CJ215 staining **h)** Fluorescence intensity in CJ215 stained cells treated with erastin and staurosporine with or without addition of Annexin V proteins show decreased fluorescence upon Annexin V blocking,. N=5 pooled from 3 independent experiments. Mean ± s.d. p-values are reported above the lines (Ordinary one-way ANOVA with Tukey’s multiple comparisons).

To further characterize this phosphatidyl serine labeling in ferroptotic cells, we utilized a commercially available dye called AnnexinVivo (which is distinct from the Annevin V protein for Annexin V staining) from PerkinElmer that is also suitable for apoptosis staining in vivo according to the manufacturer. Recent studies in our lab also identified CJ215^26^, as a novel pan-cancer agent that is chemically derived from indocyanine green (ICG) and offers a unique opportunity for cancer targeting. Our previous studies revealed that unlike ICG, which accumulates into tumors via the Enhanced Permeability and Retention (EPR) effect, CJ215 forms complexes with serum proteins^26^ upon in vitro incubation or in vivo injection reminiscent of other tumor-seeking carbocyanine dyes^27,28^. These dye-protein complexes are taken up avidly by tissues with high metabolism such as tumors or immune cells as energy sources, leading to high tumor-to-muscle ratio uptake – similar to the uptake of glucose in tumors. Our previous studies also demonstrated that CJ215 could bind to cellular membranes undergoing apoptosis and can be used to label wounds where cell death and platelet activation occurs^26^. This led us to speculate whether CJ215 can also be used as an imaging agent for ferroptosis. To ascertain whether AnnexinVivo and CJ215 can be used to image cell death, we tested the labeling of AnnexinVivo, CJ215 and ICG in cells treated with ferroptosis inducers Erastin, RSL3 and apoptosis inducer Staurosporine. We treated cells with erastin and staurosporine for 24 hours or with RSL3 for 6 hours and added the AnnexinVivo, CJ215 or ICG in the last 6 hours of this incubation period. Following incubation and washing, cells were imaged with the Licor Odyssey system for quantification. Our results showed ICG levels did not increase in any of the cells treated with either ferroptosis or apoptosis inducers, indicating the loss of membrane structural integrity during ferroptosis does not lead to increased dye uptake on its own unless dye molecules have a target, they can bind to in order to avoid being washed out and remain within the cells during the wash step. In contrast both AnnexinVivo and CJ215 can be used for monitoring RSL3 and Erastin induced ferroptosis in vitro, while CJ215 can also be used as an apoptosis imaging agent **(Figure 2b**). These results are also consistent with our findings with HT1080 cell lines treated with the same dyes and cell death inducers **(Supp Fig 4)**. It is also important to note since there are gaps in lipid membranes as well as phosphatidyl serine exposure in ferroptosis, there are more targets for AnnexinVivo and CJ215 to bind, making them more effective labels for ferroptosis. Overall, our results indicate CJ215 can serve as an ideal imaging agent for monitoring both ferroptosis and apoptosis similar to state-of-the-art imaging solutions.

Next, we tested whether ferroptosis inhibition can be monitored using CJ215 and AnnexinVivo dyes. Our results showed that erastin treatment of MDA-MB-435 cells led to increased AnnexinVivo binding, which can be inhibited with several novel ferroptosis inhibitors identified in this study, as well as those previously known ^1^ **(Figure 2c)**. In contrast, AnnexinVivo did not show an increase in HT1080 cells treated with erastin **(Supp. Fig. 5a)**, but did show an increase in HT1080 cells treated with RSL3 **(Supp. Fig. 5b)**. These results suggest that AnnexinVivo can serve as an important ferroptosis labeling agent, but it may show different characteristics depending on the cell type such as MDA-MB-435 and HT-1080 cell lines we used in this study. Furthermore, we tested whether erastin treatment increases CJ215 binding and observed an increase in dye binding upon ferroptosis induction **(Figure 2d and f)** which can be prevented with ferroptosis inhibitors. RSL3-induced ferroptosis also led to an increase in CJ215 accumulation in the cells which was in turn reduced by ferroptosis inhibitors **(Figure 2e and g)**. CJ215 staining also functioned in HT1080 cells with both ferroptosis inducers erastin and RSL3 **(Supp. Fig. 6 a and b)** indicating the flexibility of this dye over current commercial solutions. In order to show CJ215 also binds to phosphatidyl serine species, we performed a blocking assay where we compared CJ215 staining of erastin induced ferroptosis and staurosporine induced apoptosis in the presence and absence of unlabeled Annexin V protein (distinct from Annexin V staining) to block phosphatidyl serine binding. Our results showed CJ215 labeling of the erastin and staurosporine treated cells was decreased upon Annexin V addition, indicating phosphatidyl serine binding is one of the mechanisms contributing to CJ215 retention **(Figure 2h)**.

Overall, our previous publications on lipid membrane binding of CJ215 in cells undergoing apoptosis^26^ along with our current findings on reducing the cell labeling with annexin V blocking experiments indicate phosphatidyl serine is a target for CJ215 dyes. It is important though to make the distinction that while this labeling occurs in apoptosis only on the outer membrane leaflet through exposure (flipping) ofphosphatidyl serine exposure on the outer leaflet, in the case of ferroptosis, dyes can enter into the cells through nanosized gaps in the membrane induced by ferroptosis and also bind to phosphatidylserine in the inner leaflet of the membrane or on mitochondria and endoplasmic reticulum. Yet, CJ215 derivative ICG did not lead to an increased accumulation in cells treated with ferroptosis inducers which discards the possibility that increased CJ215 staining is only due to nonspecific uptake of the dye across damaged membranes. The most likely mechanism of CJ215 labeling in ferroptotic cells is utilization of gaps on the membrane to bind phosphatidyl serine molecules (or, of course, additional unknown targets) on the outer *and* inner leaflets. Collectively, these results show that CJ215 uses unique mechanisms in apoptosis and ferroptosis to bind to cellular structures and gets retained on the cell membrane, which can be utilized to image the effect of apoptosis or ferroptosis inducers and their inhibitors.

### In vivo imaging of ferroptosis based therapies

At the moment, there is no in vivo imaging agents that can monitor ferroptosis therapy. This need led us to test both AnnexinVivo and CJ215 in vivo to assess their tumor labeling and their ability to monitor ferroptosis treatment response in mice upon therapy with ferroptosis inducers. To understand general tumor targeting capabilities of these imaging agents, we generated subcutaneous xenograft HT1080 models in the flank region of the mice which were then injected with either AnnexinVivo or CJ215 without inducing ferroptosis to determine dye biodistribution. Our results showed that 24 hrs post injection Annexin Vivo had non-specific uptake in kidneys and other organs and was retained in these areas throughout the 3 day imaging period. Crucially, AnnexinVivo showed no tumor uptake **(Figure 3a, Supp. Fig. 7a-c)**. In contrast, as shown previously by us CJ215^26^ readily accumulated in tumor regions within 24 hours, followed by gradual clearance from other organs while being retained in the tumor site **(Figure 3b, Supp. Fig. 7d-f)**. These results collectively show the power of CJ215 over AnnexinVivo in tumor labeling.

**Figure 3.**
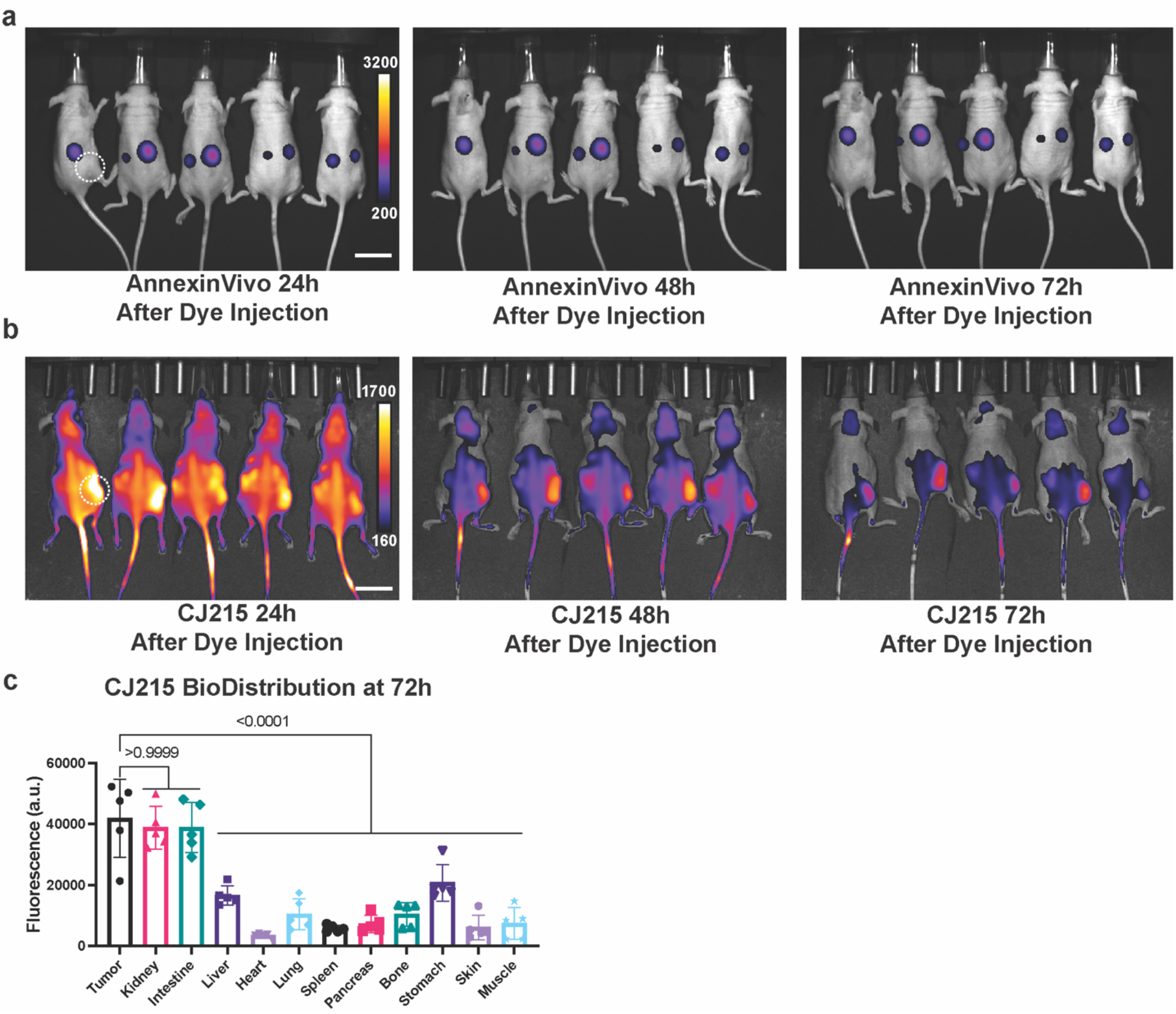
CJ215 offers enhanced targeting and contrast in vivo for tumor labeling. **a)** IVIS imaging of HT1080 xenograft tumor bearing nude mice injected with AnnexinVivo dye shows kidney localization at 24, 48 and 72 hours, **b)** IVIS imaging of HT1080 xenograft tumor bearing nude mice injected with CJ215 dye shows tumor localization with high background at 24, 48 and 72 hours, **c)** Biodistribution of CJ215 at different organs of HT1080 xenograft tumor bearing nude mice at 72 hours show increased fluorescence in tumor, intestine and kidney showing the excretion routes of CJ215, n=5 mice. Mean ± s.d. p-values are reported above the lines (Ordinary one-way ANOVA with Tukey’s multiple comparisons). Scalebar 50 mm.

While CJ215 had increased uptake in tumor sites compared to AnnexinVivo, another consideration was the rate of tumor uptake and clearance. Our results showed HT1080 xenograft tumors had a lower Contrast-to-Noise Ratio (CNR) that reached an appreciable level only around 72 hours after dye injection **(Figure 3c, Supp. Fig. 8)**. Given ferroptosis typically occurs within the 24-48 hour range after erastin addition, and dye accumulates in cells already undergoing ferroptosis, this slow accumulation and late CNR increase in HT1080 tumors prevented analysis in the time of interest. To properly image tumor responses with increased CNR at the correct imaging window, we wanted to employ an alternative and orthotopic xenograft model that achieves a high CNR faster, which led us to utilize MDA-MB-435 cell lines injected into mammary fat pads. In this study, we injected version of erastin optimized for in vivo use, Imidazole Ketone Erastin(IKE)29 either alone, or in combination with ferroptosis inhibitor liproxstatin, and waited 24 hours for the drug to reach the tumor site and induce ferroptosis. After this initial 24 hours, we injected CJ215 to the IKE treated, IKE+Liproxstatin treated and the control groups, and imaged the mice in 24-hour intervals until 72 hours post dye injection **(Figure 4a)**. At the end of the imaging experiment, we sacrificed the mice to study dye uptake and clearance routes.

**Figure 4.**
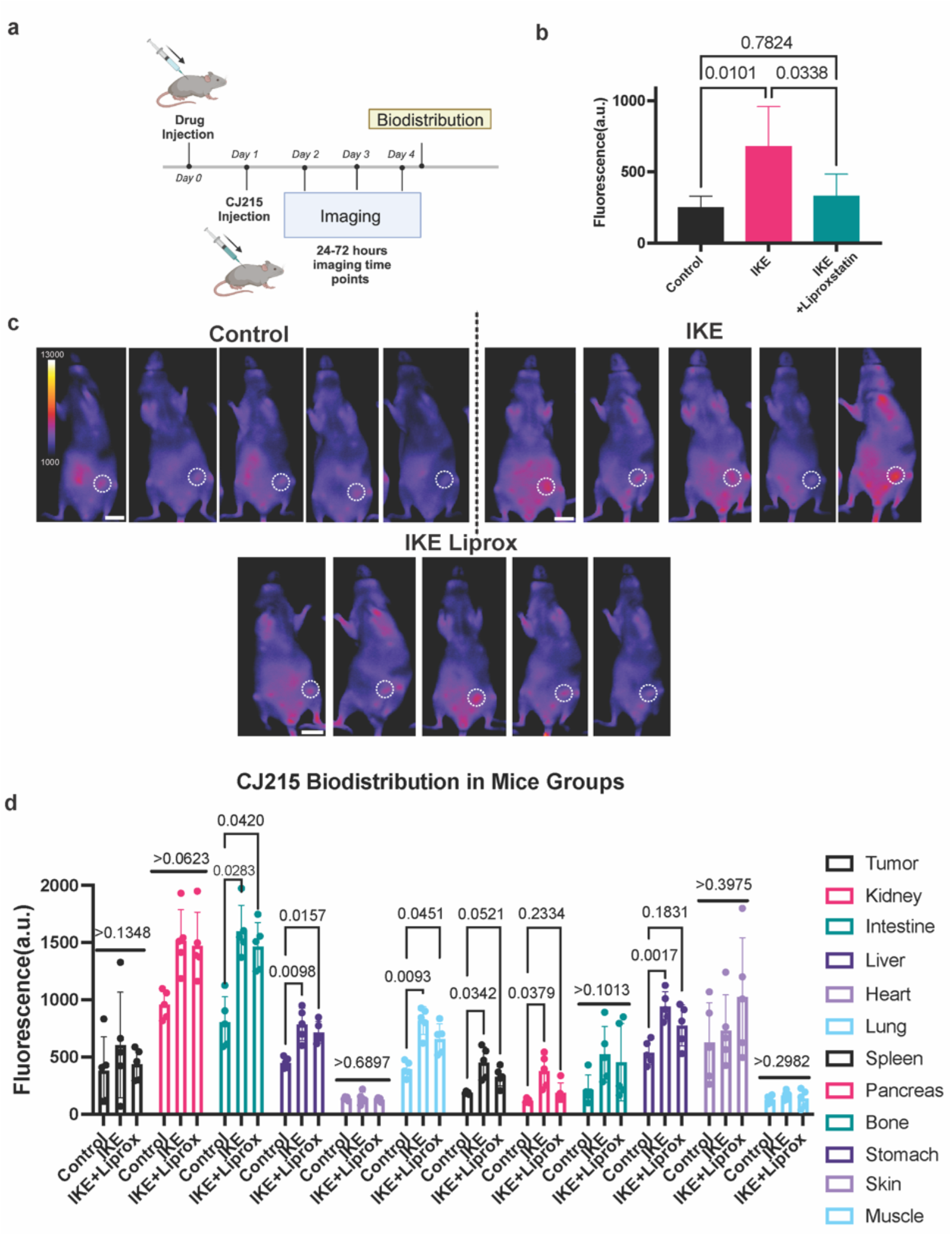
In vivo CJ215 can detect ferroptosis in vivo for monitoring ferroptosis therapies or side effects of ferroptosis drugs. **a)** Scheme outlining the experimental workflow for ferroptosis imaging from the IKE injection to imaging and biodistribution studies., **b)** Quantification of tumor uptake at 24 hours after dye injection by substracting the background CJ215 accumulation in each mice across control, IKE treated and IKE-Liproxstatin treated groups. n=5 mice. **c)** Comparison of Contrast-to-Noise Ratio of CJ215 imaging in tumor bearing mice treated with vehicle (control group), IKE and IKE-Liproxstatin imaged 24 hours after dye injection **d)** Quantification of CJ215 fluorescence in different organs indicating ferroptosis levels in mice groups treated with vehicle(control), IKE and IKE-Liproxstatin indicating off-target effects of IKE treatment in inducing ferroptosis in specific organs,Mean ± s.d. p-values are reported over the lines (Ordinary one-way ANOVA with Tukey’s multiple comparisons). Scalebar 20 mm.

Our results revealed in the case of Annexin Vivo imaging with HT1080 tumors, injection of IKE or rescue with liproxstatin does not increase tumor uptake as the dye premodinantly goes to kidneys. In contrast, we observed an increased retention of CJ215 in tumors of IKE treated mice compared with control mice within 24 hours of dye injection. Quantification of tumor to background ratios and images between these groups revealed a significant increase in tumor accumulation in IKE treated group, which can be inhibited by addition of ferroptosis inhibitor liproxstatin **(Figure 4b and c)**. This difference disappears at imaging time points 48 and 72 hours post dye injection where cancer cells die and accumulated CJ215 is cleared from the tumor site **(Supp. Fig. 9a-c)**. When we performed the biodistribution assays, we observed significant increases of CJ215 accumulation in IKE and IKE-Liproxstatin treated mice in specific organs such as intestine, liver and lung **(Figure 4d and Supp. Fig. 10)**. These results indicate while IKE is a potential anti-tumor agent, it can also lead to off-target effects and induce ferroptosis in other organ types which rely on Systems Xc^-^ as a part of their cellular homeostasis. While liproxstatin co-injection significantly decreased most of these organ accumulations (with the exception of intestines), it was not sufficient to protect the entire mouse. Overall, these results indicate, beyond tumor imaging, CJ215 accumulation also offers unique opportunities to test ferroptosis in physiological settings and can be employed to study side-effects of ferroptosis inducing drugs or effectiveness of ferroptosis inhibitors.

## Conclusion

In this study, we explored how ferroptotic cell death is accompanied by changes in membrane dynamics and how lipid peroxidation accumulation results in membrane damage which we characterized with different sizes of quantum dots and various dyes, taking advantage of their size dependent emission spectra changes and binding characteristics. Our results showed membrane damage opens nano-sized gaps using a machinery similar to platelet activation. We also showed many platelet activation inhibitors in PIP2-PLC-IP3-PKC pathways can also be utilized as ferroptosis inhibitors. It is interesting to note how platelet activation is also implicated in stroke and plaque formation, which are shown to have underlying ferroptotic components^30^. These similarities between platelet activation and ferroptosis can also explain why ferroptosis is not an immunogenic cell death despite a loss of membrane integrity unlike necroptosis^31^. We have further shown how ferroptosis induced nano-sized cell membrane gaps during late stage ferroptosis can lead to Annexin V binding either through phosphatidyl serine exposure or through traversing cell membrane and binding to phosphatidyl serine molecules on the inner leaflet. We reasoned phosphatidyl serine targeting approaches such as Annexin V based assays would serve as potential ferroptosis labeling agents, and we demonstrated this using both conventional Annexin V stain as well as a commercially available in vivo imaging agent called AnnexinVivo. Despite their use in the in vitro setting, annexin based imaging agents were not highly successful in detecting apoptotic cells in vivo in the past^32^, and our experience with AnnexinVivo revealed non-specific uptake by other organs and limited tumor labeling. Therefore, we turned to a novel carbocyanine dye, CJ215, that binds to serum proteins both in vitro and in vivo to act as a transient passenger for tumor uptake. Given the excess nutrient demands of tumors compared with the rest of the body^33^ these dye-protein complexes preferentially accumulate in the tumor sites. In the ferroptotic and apoptotic cancer cells, however, CJ215 show even a more pronounced accumulation which can be prevented using unlabeled Annexin V protein as a blocking agent. It is important to note these two mechanisms are not mutually exclusive and can work in tandem where dye-albumin complexes are essential for bringing the CJ215 into the tumor site and phosphatidyl serine binding can lead to increased retention on cancer cells. Overall result would be high CNR in tumors undergoing cell death which can be utilized for therapy monitoring.

In vivo application of CJ215 still suffers from several challenges given it only binds to the cells during the late stage of ferroptosis. This requires a comprehensive knowledge of biodistribution of ferroptosis inducers, the response time of tumors to the ferroptosis inducers, as well as dye uptake and clearance, all of which would contribute to the success of CJ215 as a therapy monitoring dye. While our efforts were successful to image changes in tumors undergoing ferroptosis in MDA-MB-435 tumors, HT1080 xenografts had slow clearance which increased the background and lowered tumor selective Contrast-to-Noise Ratios. Thus, future applications of CJ215 would require an in-depth knowledge of tumor type, ferroptosis inducer and imaging agent, and their successful coordination for an optimized protocol that is unique for each situation. Finally, we are aware phosphatidyl serine exposure and membrane defects are not unique biomarkers for ferroptosis, and we do not imply CJ215 imaging can be used to prove a drug is in a fact inducing ferroptosis. Yet, we have established a ferroptosis imaging approach that is applicable to an in vivo setting which was greatly needed. Here, a known ferroptosis inducer, combined with a suitable rescue experiment via ferroptosis inhibitors such as liproxstatin can offer unique insights into in vivo ferroptosis biology. We believe CJ215 based imaging approaches have wide-ranging applications not only for tumor labeling, but also reporting side effects of ferroptosis inducers in diverse organ systems along with a cancer therapy monitoring approach for both apoptosis and ferroptosis based treatment strategies.

## Acknowledgments

We would like to thank ProImaging for providing CJ215 for this investigation and the small animal imaging core at MSKCC, as well as RARC at MSKCC for their guidance in animal experimentation and husbandry.

## Materials and Methods

### Cell Culture

MDA-MB-435 and HT1080 cells were obtained from American Type Culture Collection(ATCC). MDA-MB-435 cell line was cultured in RPMI 1640 Medium(Thermo Fisher), 10 % Fetal Bovine Serum (FBS) and 1% Penicilin-Streptomycin(Pen-Strep) Solution. HT1080 cell line was cultured with DMEM medium (Thermo Fisher) with 10% FBS and 1% Pen-Strep. Media was changed in every 3 days to ensure nutrient availability and cells were passaged at 80% confluence with 0.05% Trypsin EDTA (Thermo Fisher). Cell viability and health were constantly monitored during the course of the study.

### Quantum Dot Uptake

Cultured cells were washed with Phosphate Buffer Saline(PBS) and trypsinized. Cells were centrifuged to remove excess trypsin and were resuspended in their respective culture media. Cells were placed in 24-well plates with 75,000 cells per well with a media volume of 500 μL. The cells were incubated in 37 °C with 5% CO^2^ levels overnight for surface attachment. Each well was treated with either vehicle or erastin(10 μM) (Cayman Chemicals) for 6 or 18 hours. After the incubation, the cells were washed with PBS and trypsinized for flow cytometry preparation. They were subsequently centrifuged for 5 minutes at 1000 rpm and resuspended in PBS containing 2% FBS. Cells were centrifuged again and resuspended again in PBS-2%FBS. Then, the cells were incubated with DAPI (1 μM), carboxyl coated QD450, QD550, QD650 quantum dots(Sigma Aldrich) (1 μL from the stock solution) separately for 5 minutes on ice and were analyzed in flow cytometry(MACS Quant). Given the distinct spectral nature of quantum dots and dyes, we could distinguish each quantum dot type in different channels. We followed the 5 minute incubation protocol on ice to prevent non-specific uptake and employed same surface coated Quantum dot types to ensure only size remains a factor in quantum dot uptake. We used the negatively charged carboxyl coated quantum dots to prevent non-specific charge based interactions with the cell membrane. We also normalized the QD uptake to the vehicle treated cells to ensure nonspecific cell surface binding is accounted for. The change in cellular fluorescence was analyzed using FlowJo program. Quantum dot sizes were also confirmed using Malvern PanAnalytical Zeta Sizer measurements.

### Cytotoxicity

Cells were trypsinized and seeded on 96-well plate with 10,000 cells per plate in 150 μL. The cells were incubated in 37 °C with 5% CO_2_ levels overnight for surface attachment. Only the center 60-wells were used for the assays to prevent the edge-effects. The cells were treated with Erastin(10 μM) or RSL3(2 μM)(Cayman Chemicals) in the presence and absence of the inhibitors and activators. 2-APB, D609, SF1670 and Rotenone were also purchased from Cayman Chemicals and utilized with final concentrations of 2 μM, 5 μM, 0.2 μM and 1 μM respectively. The concentrations of the other chemicals are given in Supplementary Tables all of which was purchased from MedChem. Cells were treated with ferroptosis inducers and inhibitors for 24 hours and cytotoxicity was measured using Cell Titer Glo2 Assay (Promega) based on the manufacterer’s instructions. The cells were incubated with Cell Titer Glo2 reagent for 10 minutes and this mixture is transferred to a black-walled plate to prevent cross-well light detection. The bioluminescence was measured using Molecular Devices SpectraMax iD5 Multi-Mode Microplate Reader with 1000 ms integration time with open spectral detection.

### Mitochondrial Membrane Potential

Cells were trypsinized and seeded on 96-well plate with 20,000 cells per plate in 150 μL. The cells were incubated in 37 °C with 5% CO_2_ levels overnight for surface attachment. Only the center 60-wells were used for the assays to prevent the edge-effects. The cells were treated with Erastin(10 μM) in the presence and absence of the inhibitors for 6 hours. The cells were then incubated with JC10(4 μM) for 30 minutes and washed with PBS twice. The increase in fluorescence was measured using Molecular Devices SpectraMax iD5 Multi-Mode Microplate Reader with 560 nm excitation and 600 nm emission.

### Lipid Peroxidation

Trypsinized cells were placed in 24-well plates with 75,000 cells per well with a volume of 500 μL. The cells were incubated in 37 °C with 5% CO_2_ levels overnight for surface attachment. Each well was treated with erastin(20 μM) for 6 hours either with our without the ferroptosis inhibitors. For the last 30 minutes of the incubation, C11-BODIPY^581/591^ (Molecular Probes) (2 μM) dye was added to cell culture media based on manufacturer’s instructions. After the 30 minute incubation with the dye, the cells were washed with PBS and trypsinized for flow cytometry preparation. They were subsequently centrifuged for 5 minutes at 1000 rpm and resuspended in PBS containing 2% FBS. Later, they were centrifuged again and resuspended in PBS-2%FBS containing DAPI (1 μM) for 5 minutes and they were analyzed using MACS Quant flow cytometry. C11 Bodipy dye works by increasing its green fluorescence in the presence of lipid peroxidation and we used vehicle treated wells as a baseline to measure relative increase in lipid peroxidation levels. The change in cellular fluorescence was analyzed using FlowJo program.

### Annexin V staining

Trypsinized cells were placed in 24-well plates with 75,000 cells per well with a volume of 500 μL. The cells were incubated in 37 °C with 5% CO_2_ levels overnight for surface attachment. The cells were treated with Erastin(10 μM) for 6 or 18 hours or RSL3(2 μM)(Cayman Chemicals) for 2 hours in the presence and absence of Liproxstatin (1 μM). After the incubation, the cells were washed with PBS and trypsinized for flow cytometry preparation. They were subsequently centrifuged for 5 minutes at 1000 rpm and resuspended in PBS containing 2% FBS. Cells were centrifuged again and resuspended again in PBS-2%FBS. Then the cells were incubated for 20 minutes with Annexin V-FITC stain with PI dye to measure membrane integrity (MedChem) based on manufacturer’s instuctions. The cells were analyzed using MACS Quant flow cytometry. Annexin V-FITC binding indicates either phosphatidyl serine exposure or influx of Annexin V into the cell via nano-sized gaps on the cell membrane surface. The change in cellular fluorescence was analyzed using FlowJo program.

### Annevin Vivo and CJ215 Imaging In vitro

Trypsinized cells were placed in 24-well plates with 75,000 cells per well with a volume of 500 μL. The cells were incubated in 37 °C with 5% CO_2_ levels overnight for surface attachment. The cells were treated with Staurosporine(2 μM), Erastin(10 μM) for 18 hours or RSL3(1 μM) for 6 hours in the presence and absence of ferroptosis inhibitors. CJ215 (0.645 mM), Annexin Vivo (1:1000 dilution from stock solution) and ICG(0.645 mM), dyes were added to the cell culture media, incubated for 6 hours and washed with PBS. For Annexin V blocking experiments cells were co-incubated with CJ215 and unlabeled Annexin V proteins(Abcam)(2 μg/well). The cells were imaged using Licor Odyssey CLX imager in both 680 and 800 nm channels and images were quantified using ImageJ.

### Annexin Vivo and CJ215 Imaging In vivo

All mouse handling, experimentation, imaging, and housing was performed according to NIH guidelines and via approved IACUC protocols at MSKCC. Tumor xenografts were developed using 5.0×10^6^ HT1080 cells were resuspended in 100 µL Matrigel-media mixture (1:1 ratio) and subcutaneously injected into flank of female FoxN1^nu^ mice (6-8 weeks). Tumor growth was monitored visually and dye injection was performed when tumors exceeded 50 mm^3^. Mice received a 2mg/kg intravenous injection of CJ215 suspended in clinical grade dextrose or 100 µL AnnexinVivo solution per mice as described by manufacturer’s instructions. For the therapy imaging, mice were injected with vehicle(70% PBS, 20%PEG400, 10% ethanol), IKE alone(20 mg/kg), or IKE(20 mg/kg) with Liproxstatin(10 mg/kg) 24 hours before the dye injection. After the CJ215 injection(2 mg/kg), we waited another 24 hour for dye clearance to achieve an optimal clearance and high signal-to-noise ratio in the tumor region. Mice were imaged using IVIS-CT with 2 s exposure time and f# of 1 with small binning using ICG filter sets(745 excitation and 840 nm emission) every 24 hours. During this time mice was under isofluorane anesthesia at an induction of 3% v/v and anesthesia was maintained of 1-2% v/v in O_2_ where mice were kept on a heating pad during the course of imaging experiments. After the imaging sessions, mice were euthanized with CO_2_ and their organs were harvested. Biodistribution of the dye was assessed by isolating the organs: tumor, liver, kidneys, hearts lungs, spleen, pancreas, bones, stomach, intestines, skin, and muscle. We performed the same imaging conditions for organ imaging, exported the tiff files that contain the fluorescence levels and analyzed the images using ImageJ/Fiji with ROIs drawn in background regions on live mice, tumor regions and individual organs. When we processed the live mice imaging experiments, we determined the background of each mice with measuring the relative dye accumulation in their chest cavity and subtracted half of this value from the total image to achieve a consistent background for the visualization. We analyzed the results using GraphPad Prism software where we calculated the means and standard deviations of each group and performed one-way Anova with multiple comparisons or t-tests where applicable as detailed in each figure legend.

