## Supplementary Information for "In vivo imaging of ferroptosis through nanodynamic changes in lipid membranes"

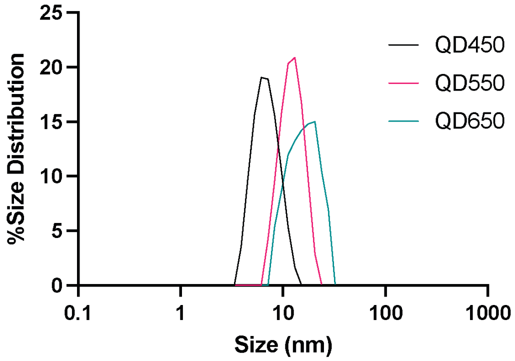


**Supplementary Figure 1. Size distribution of quantum dots used in the study** ZetaSizer measurements of QD450, QD550 and QD650 showing differential size distribution

**a**

| Name of the Inhibitors and Activators | Effect of the Chemicals | Concentration |
|---|---|---|
| Isoprenaline | β-adrenergic receptor agonist | 5 μM |
| Treprostinil | DP1 and EP2 agonist | 5 μM |
| Laropiprant | DP receptor antagonist | 5 μM |
| RO1138452 | IP (prostacyclin) receptor antagonist | 5 μM |
| Asapiprant | $DP_1$ receptor antagonist | 5 μM |
| Grapiprant | EP4 receptor antagonist | 5 μM |
| Misoprostol | Synthetic analogue of prostaglandin E1 (PGE1) | 5 μM |
| Iloprost | Prostacyclin (PGI2) analogue | 5 μM |
| Seratrodast | Thromboxane A2 receptor (TP) antagonist | 5 μM |

Table 1. List of G-Protein Coupled Receptor Modulators for Adrenergic, Prostaglandin and Thromboxane Receptors and their Derivatives

**b**

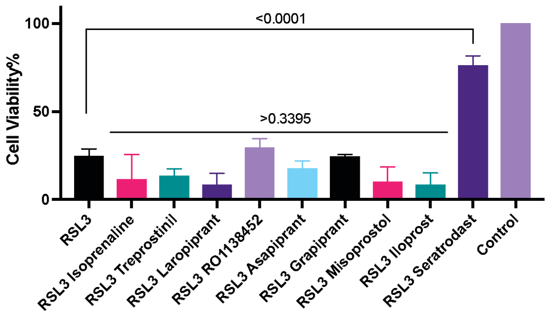


**Supplementary Figure 2. Effect and concentration of different GPCR modulators tested in the study and their effect on RSL3 induced ferroptosis a)** Table showing the effect and concentration of different GPCR inhibitors used in this study **b)** Cytotoxicity of RSL3 induced ferroptosis treated with Adrenergic, prostaglandin and thromboxane receptor inhibitors For cytotoxicity assays n=3 biological replicates, Mean ± s.d. p-values are reported above the lines (Ordinary one-way ANOVA with Tukey’s multiple comparisons).

a

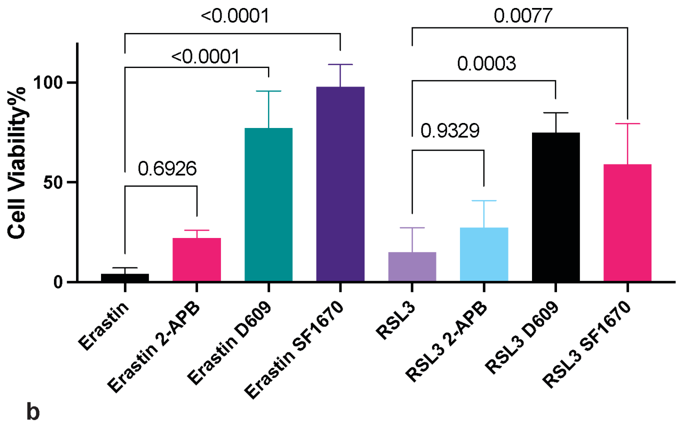


b

| Name of the Inhibitors and Activators | Effect of the Chemicals | Concentration |
|---|---|---|
| Bisindolylmaleimide I | PKCα, PKCβI, PKCβII, and PKCγ | 5 μM |
| Bisindolylmaleimide IX | PKC-α, PKCβI, PKCβII, PKC-γ, and PKC-ε | 5 μM |
| Go6983 | PKCα, PKCβI, PKCβII, PKCγ and PKCδ | 5 μM |
| Go6976 | PKCα, and PKCβI | 5 μM |

c

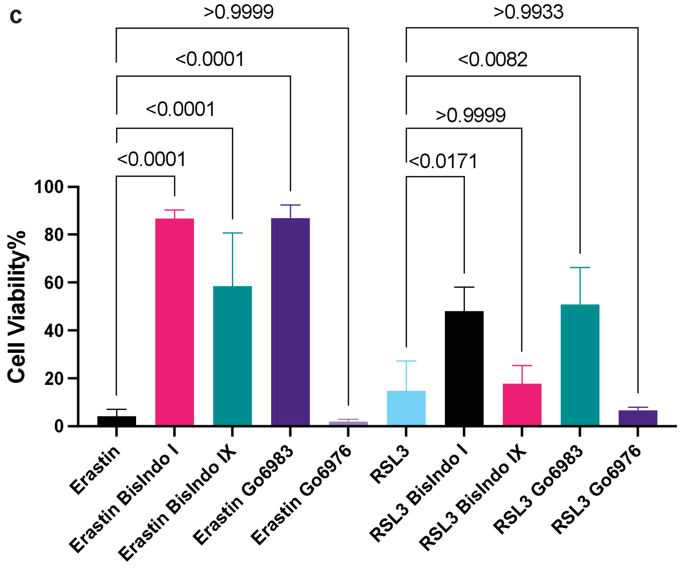


**Supplementary Figure 3. Inhibitors of Platelet activation rescues ferroptotic cell death in HT1080 cell line a)** IP3R, PLC and PTEN inhibitors rescue the cells from both erastin and RSL3 induced ferroptosis **b)** Effect and Concentration of different PKC inhibitors, **c)** Role of different PKC inhibitors and their effect on erastin and RSL3 induced ferroptosis. For cytotoxicity assays n=3 biological replicates, Mean ± s.d. p-values are reported above the lines (Ordinary one-way ANOVA with Tukey's multiple comparisons).

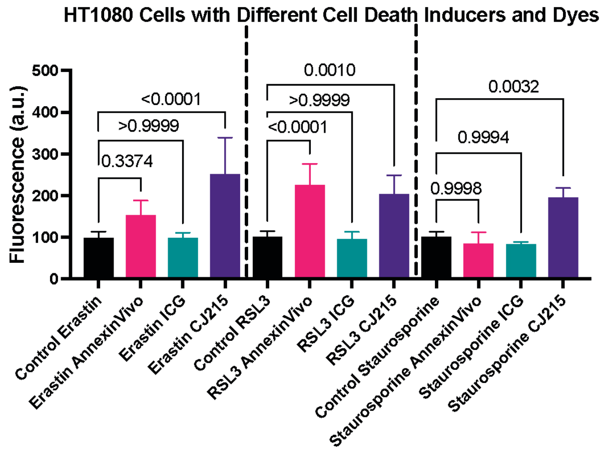


**Supplementary Figure 4. Comparing the effectiveness of different dyes in HT1080 cells upon ferroptosis and apoptosis induction** Measuring the fluorescence levels in HT1080 cells treated with erastin, RSL3 and staurosporine using three different dyes AnnexinVivo, ICG and CJ215 showing the effectiveness of CJ215 over the other contrast agents N=5 pooled from 3 independent experiments. Mean ± s.d. p-values are reported above the lines (Ordinary one-way ANOVA with Tukey's multiple comparisons).

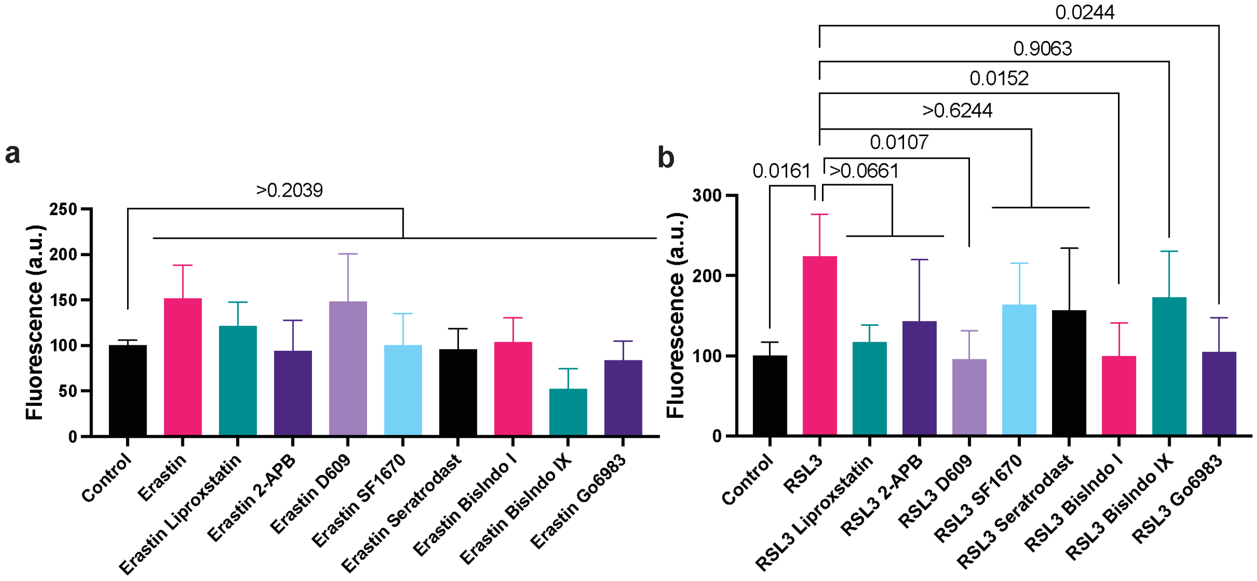


**Supplementary Figure 5. Changes in Fluorescence intensity in ferroptosis inducers and inhibitors using AnnexinVivo dyes as contrast agent in HT1080 cell line a)** Measurement of fluorescence upon 24 hours of erastin treatment (10 μM) along with ferroptosis inhibitors using AnnexinVivo staining **b)** Measurement of fluorescence upon 6 hours of RSL3 treatment (1 μM) along with ferroptosis inhibitors using AnnexinVivo staining N=5 pooled from 3 independent experiments. Mean ± s.d. p-values are reported above the lines (Ordinary one-way ANOVA with Tukey's multiple comparisons).

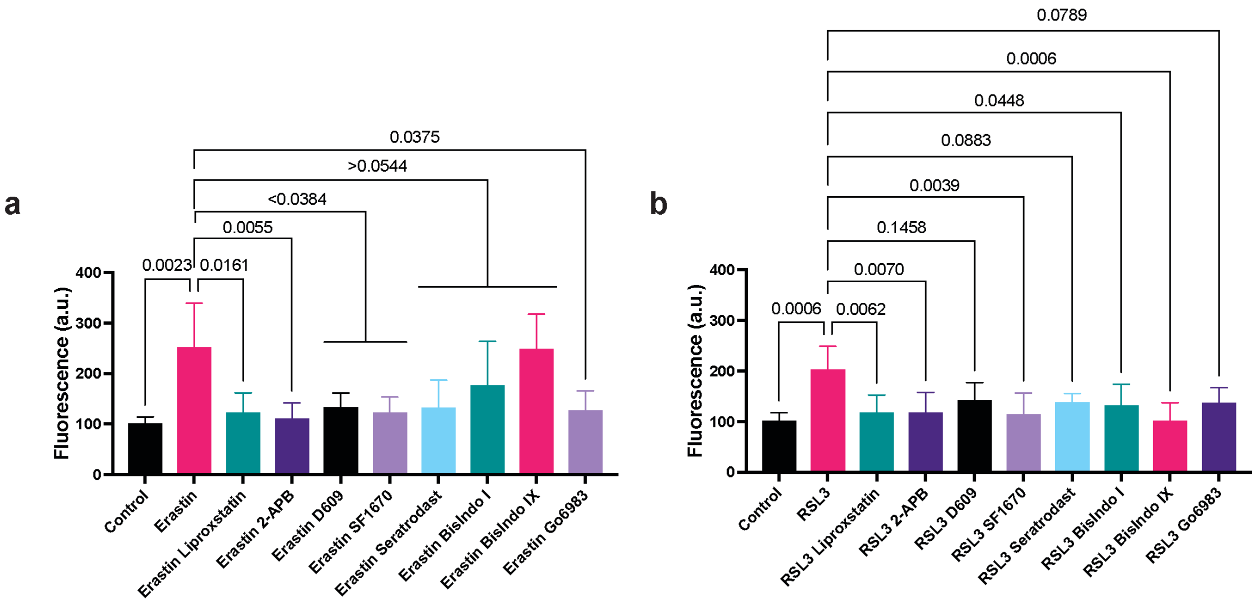


**Supplementary Figure 6. Changes in Fluorescence intensity in ferroptosis inducers and inhibitors using CJ215 dyes as contrast agent in HT1080 cell line a)** Measurement of fluorescence upon 24 hours of erastin treatment (10 μM) along with ferroptosis inhibitors using CJ215 staining **b)** Measurement of fluorescence upon 6 hours of RSL3 treatment (1 μM) along with ferroptosis inhibitors using CJ215 staining N=5 pooled from 3 independent experiments. Mean ± s.d. p-values are reported above the lines (Ordinary one-way ANOVA with Tukey's multiple comparisons).

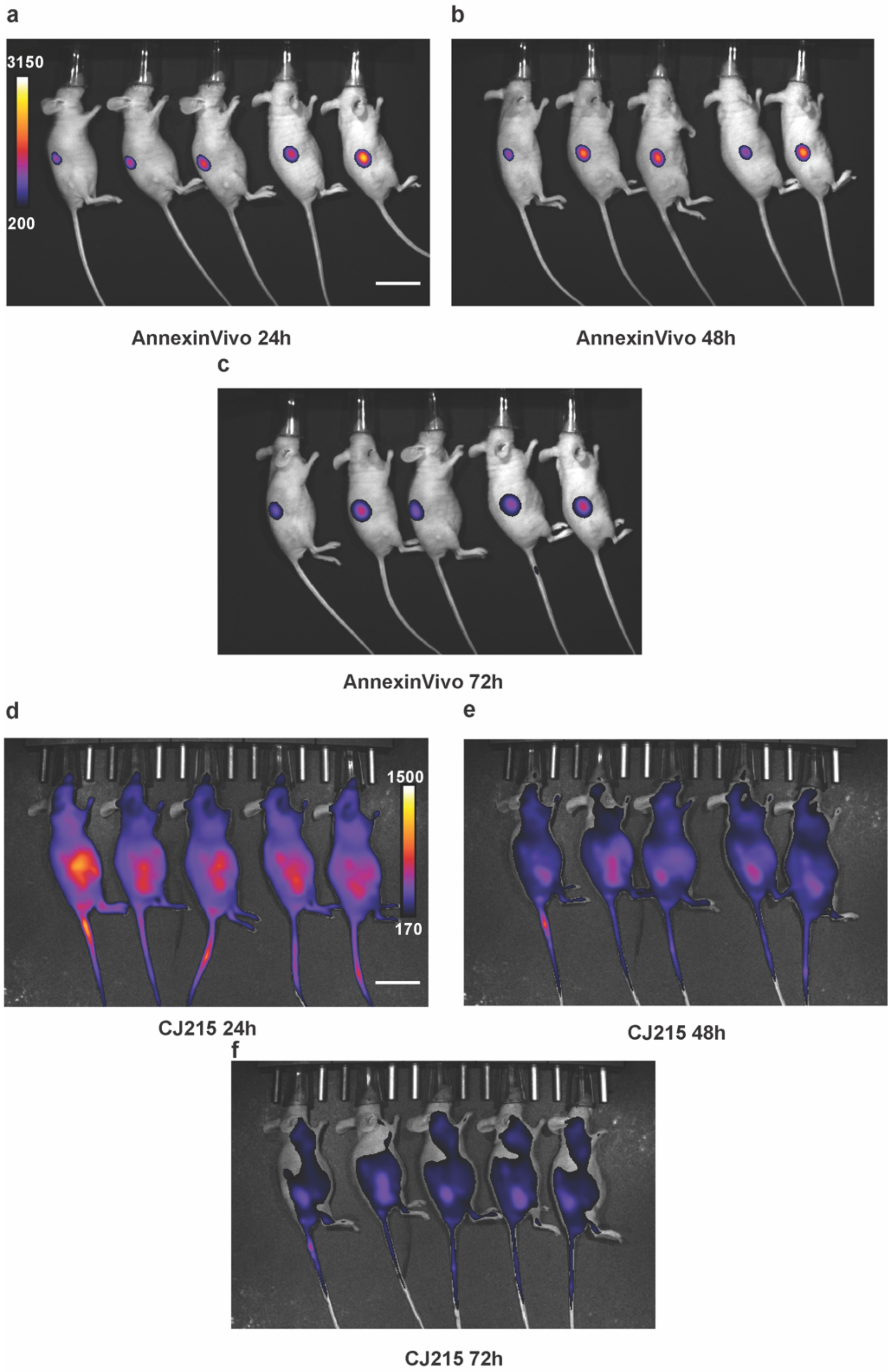


**Supplementary Figure 7. Side view of AnnexinVivo and CJ215 injected mice bearing HT1080 xenograft tumors demonstrates localization of the dyes** IVIS imaging of AnnexinVivo injected HT1080 xenograft tumor bearing nude mice at **a)** 24 hours **b)** 48 hours **c)** 72 hours at side view showing localization in kidney and not in the tumors, IVIS imaging of CJ215 injected HT1080 xenograft tumor bearing nude mice at **d)** 24 hours **e)** 48 hours **f)** 72 hours at side view showing localization in the tumor but with a high background in the rest of the body, n=5 mice. Scalebar 50 mm

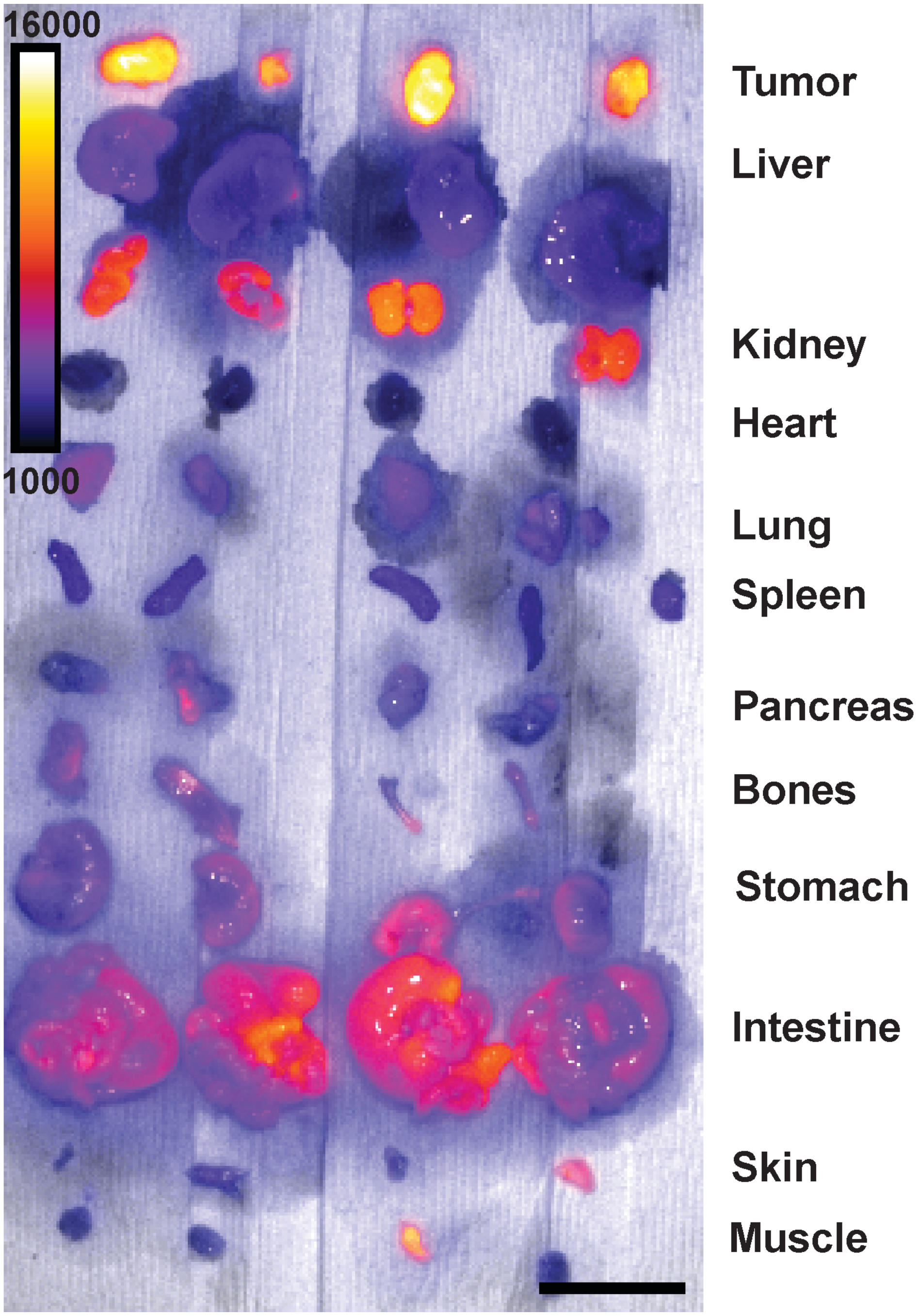


**Supplementary Figure 8. Fluorescence images of the organs in CJ215 injected mice bearing HT1080 xenograft tumors demonstrating dye biodistribution** Biodistribution of CJ215 in different organs in HT1080 xenograft tumor bearing nude mice 72 hours after injection shows kidneys are slowly excreting the dye over time, n=4 mice. Scalebar 50 mm.

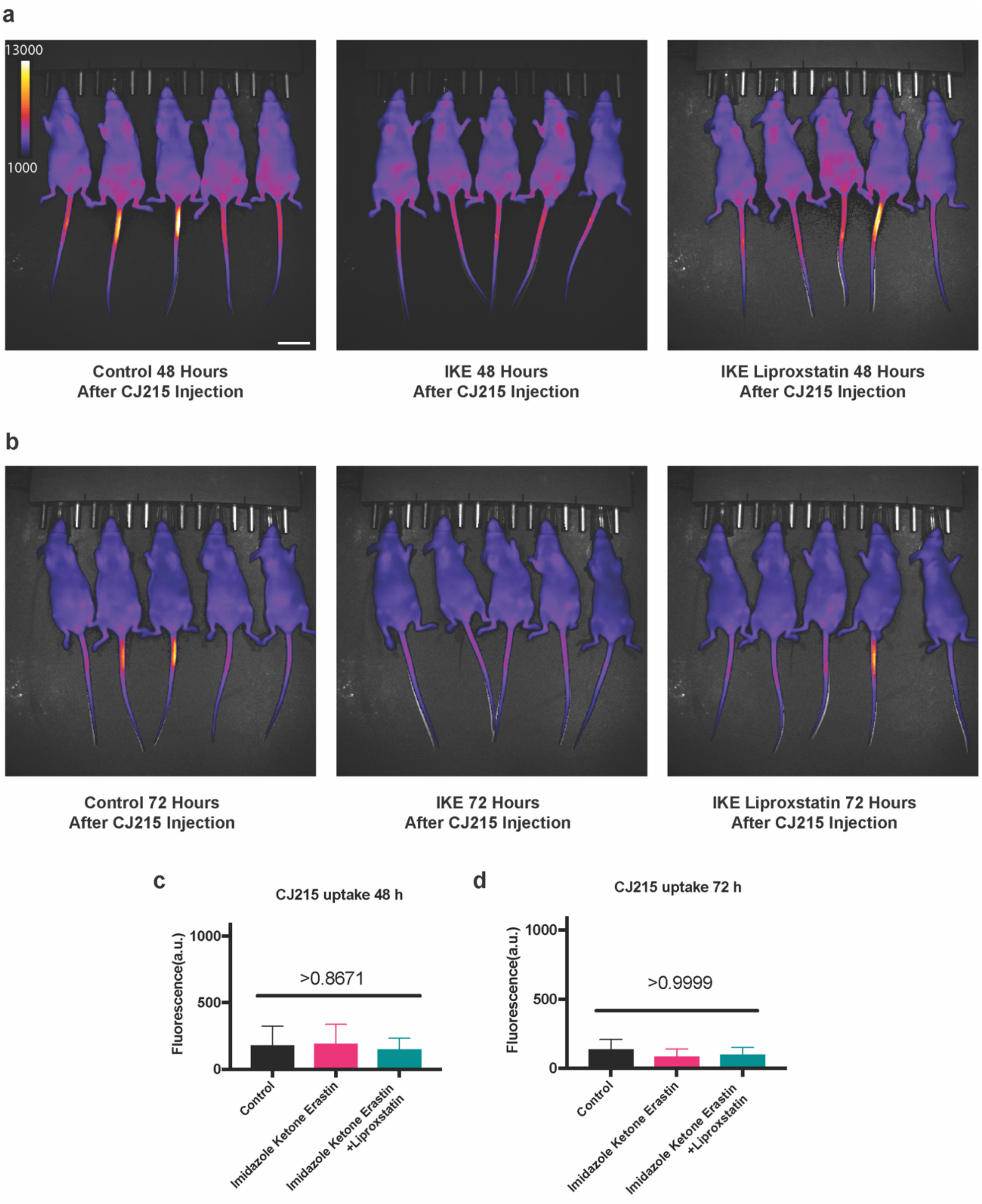


**Supplementary Figure 9. Quantification of CJ215 tumor uptake at later stages of ferroptosis therapies.** Imaging CJ215 tumor uptake in mice treated with vehicle(control), IKE and IKE-Liproxstatin at **a)** 48 hours post dye injection and **b)** 72 hours post dye injection**, c)** Quantification of tumor uptake between different mice groups reveal no difference between control and IKE treated group after 48 and 72 hours post dye injection, indicating the ferroptotic cell death is followed by clearance of the dead cells and dye. Mean ± s.d. p-values are reported above the lines (Ordinary one-way ANOVA with Tukey's multiple comparisons). Scalebar 50 mm

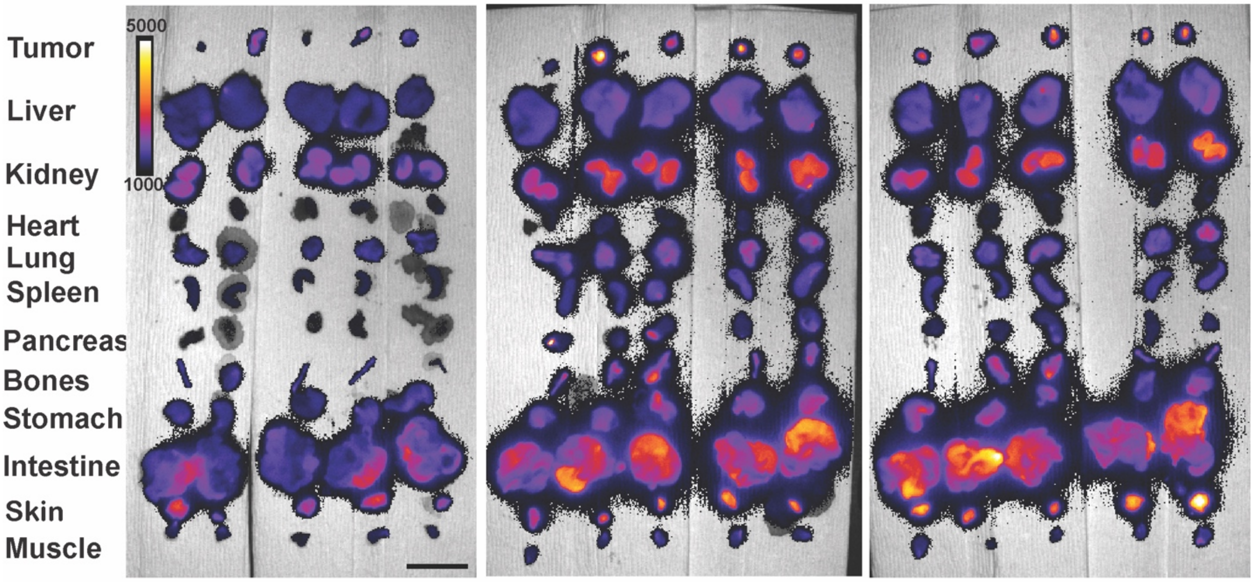


**Supplementary Figure 10. Biodistribution of CJ215 in xenograft mice models with and without ferroptosis inducers.** Biodistribution image of different organs from MDA MB 435 tumor bearing xenograft mice treated with vehicle(control), IKE and IKE-Liproxstatin, n=5 mice Scalebar 50 mm.
